# Biosurfactant polysaccharide drives dispersal of exopolysaccharide-coated myxospores in *Myxococcus xanthus*

**DOI:** 10.64898/2026.09.26.754727

**Authors:** Antoine Bignet, Nada Rezania, Ahmad A. Kezzo, Niharika Saraf, Arnaldo Nakamura, Gaurav Sharma, Frédéric J. Veyrier, Salim T. Islam

## Abstract

Bacterial spores are highly resilient cell forms, yet the organization of their protective surface layers in diderm species remains largely unexplored. For the social predatory soil bacterium *Myxococcus xanthus*, myxospore formation within multicellular fruiting body structures is a key ecological adaptation to surviving conditions of nutrient deprivation. These myxospores were long thought to be outwardly coated with so-called major spore coat (MASC) polysaccharide. Using polysaccharide-deficient mutant strains, electron microscopy, surface hydrophobicity assays, dye binding, and functional swarm complementation, we show that MASC instead forms a protective sub-surface outer-cortex layer, while the exopolysaccharide (EPS) glycocalyx inherited from vegetative cells constitutes the true outermost myxospore coat. EPS mediates inter-myxospore adhesion and supports collective behaviors, whereas secretion of biosurfactant polysaccharide (BPS) reduces myxospore surface stickiness, preventing excessive aggregation. BPS-deficient myxospores form persistent clumps, while EPS-deficient myxospores fail to aggregate or restore EPS-dependent swarm motility of vegetative cells. These results reveal a mechanistic interplay in which MASC serves as a sub-surface protective layer, EPS defines the exterior interface, and BPS tunes surface accessibility to regulate dispersal. Our findings overturn long-standing assumptions about myxospore architecture and illustrate a polysaccharide-driven strategy for controlling aggregation and readiness for germination. More broadly, these results highlight how social diderm bacteria leverage inherited and dynamically remodeled polysaccharide surfaces to coordinate multicellular development, in contrast to the rigid protein-based spore surfaces of Gram-positive species.

## INTRODUCTION

Spore formation is a widespread and vital survival strategy employed by diverse organisms to endure unfavorable environmental conditions^1^. Spores are typically characterized by metabolic dormancy, structural resilience, and long-term viability, enabling populations of these organisms to persist in the face of nutrient limitation, desiccation, high-energy radiation, and other stresses. While the structural details and developmental pathways involved vary across the multitude of spore-generating organisms, the formation of protective surface layers—often containing different polysaccharides, proteins, and lipids—is a common theme underpinning spore robustness and ecological success^1^.

In bacteria, the most widely-studied spores are those from the monoderm (“Gram-positive”) soil bacterium *Bacillus subtilis*; these endospores are produced through asymmetric cell division, generating a smaller forespore and a larger mother cell, with the latter engulfing the former, leading to successive addition of endospore layers^2^. Surrounding the *B. subtilis* endospore core (containing the chromosome, ribosomes, etc.) are thus: (i) the inner forespore membrane (a highly impermeable lipid bilayer), (ii) the germ cell wall (a thin peptidoglycan layer that becomes the functional cell wall upon germination), (iii) the cortex (a thick protective peptidoglycan layer with reduced crosslinking), (iv) the outer forespore membrane (a remnant of the engulfment process), (v) the inner/outer coat & crust (a triple-layered proteinaceous covering), and sometimes (vi) the exosporium (a loose, glycoprotein-rich layer) **(Supplementary Fig. 1)**.

Conversely, far less is known about sporulation in diderm (“Gram-negative”) bacteria. Social predatory *Myxococcus xanthus* is the premier model organism for studying complex developmental processes in diderm bacteria, including the formation of starvation-induced multicellular fruiting bodies^3^. Therein, ∼80% of aggregated vegetative cells undergo autolysis and liberate their contents while ∼10% of cells differentiate into environmentally-resistant myxospores. These dormant cell-forms share certain properties with their vegetative counterparts, while also exhibiting distinct morphological and biochemical features. Surrounding the myxospore core (containing the chromosome, ribosomes, etc.) are: (i) an inner membrane (IM) phospholipid bilayer, (ii) a periplasmic space, (iii) an outer membrane (OM) (containing lipopolysaccharide^4,5^), (iv) a “cortex” layer (of unknown composition), and (v) a compacted surface layer attributed to major spore coat (MASC) polysaccharide (formerly referred to as a coat/surface coat/extracellular coat/outer layer/slime capsule/microcyst capsule/outer sheath/cuticula)^6–13^, a view held for >63 years^14^ **(Supplementary Fig. 1)**. This surface layer is critical for conferring resistance to environmental insults and is a hallmark of both naturally-occurring and chemically-induced mature myxospores^3^.

Though MASC polysaccharide is only produced during development (for myxospores), the surface glycocalyx of vegetative *M. xanthus* cells is composed of exopolysaccharide (EPS), a specific polymer. In turn, secretion of biosurfactant polysaccharide (BPS) is required to functionally destabilize the integrity of the EPS glycocalyx, allowing for wild-type swarm spreading (mediated by Type IV pilus [T4P] extension and retraction^15,16^), single-cell motility (mediated by the motorized trans-envelope Agl–Glt–CglB gliding apparatus^17,18^), and tolerance to toxic compounds^19–21^. Each of MASC, EPS, and BPS is assembled-and-secreted by a separate pathway^22^. Within *M. xanthus* communities, both EPS and BPS act as “shared goods” that can complement respective deficiencies when supplied *in trans*^19,20,23^.

To form fruiting bodies under nutrient-poor conditions, *M. xanthus* enters into a developmental cycle in which vegetative cells aggregate to form mounds, inside of which the above-described autolysis and myxospore differentiation takes place. The aggregation of myxospores within fruiting bodies may facilitate spore survival and/or synchronized germination to facilitate group feeding. While aggregation of cells and/or spores can arise from both active and passive mechanisms—including physical entrapment, cell-surface interactions, and extracellular matrix components—its molecular basis remains incompletely understood. Surface properties such as charge, hydrophobicity, and polysaccharide composition are key determinants of passive cell–cell interactions, but in *M. xanthus*, these are complicated by the production of BPS that can modulate cell aggregation dynamics.

To better understand the intrinsic aggregative properties of myxospores, it is essential to investigate their surface characteristics in the absence of biosurfactant influence. This approach allows us to isolate the contributions of the spore envelope itself—particularly the MASC polysaccharide layer—to aggregation phenomena, without the confounding effects of extracellular surfactants. In this study, we examine the aggregation behavior of *M. xanthus* myxospores under biosurfactant-free conditions, aiming to elucidate the role of spore surface composition and architecture in passive aggregation. Our findings provide insight into the biophysical principles underlying spore clustering and offer a clearer picture of how surface specialization supports dormancy and survival in this model developmental system.

## RESULTS

### BPS-and MASC-pathway expression is upregulated during late development

Given the pivotal effects of BPS on the EPS of vegetative *M. xanthus* cells and the ensuing physiological changes^19–21^, we sought to probe the importance of secreted polysaccharides on cells undergoing development. We first examined RNAseq data for vegetative *M. xanthus* cells that had been spotted on a nutrient-poor agar substratum to induce sporulation and fruiting body formation^24^. Specifically, the expression levels of the outer-membrane polysaccharide export (OPX) genes *wzaB/S/X* (corresponding to the BPS/MASC/EPS pathways) were followed over the course of development. The encoded OPX proteins are essential periplasmic components of each pathway as they link the respective integral inner-and outer-membrane machinery to mediate export of internally-produced polysaccharide to the cell exterior^22,25,26^. Using this readout, the MASC pathway levels were found to be basal until late in development at which point they were highly upregulated **(Fig. 1)**, consistent with previously-published reports^27,28^. Intriguingly, expression of the BPS pathway was also found to increase during the same later stages of development as those shown for increased MASC pathway production, suggesting a possible role for BPS during sporulation **(Fig. 1)**. Unexpectedly, EPS-pathway levels did not decrease at all throughout the life cycle, even during later developmental stages at which sporulation is taking place **(Fig. 1)**. Taken together, these data heavily implicate each of BPS, MASC, and EPS as being important during various stages of development.

**Figure 1.**
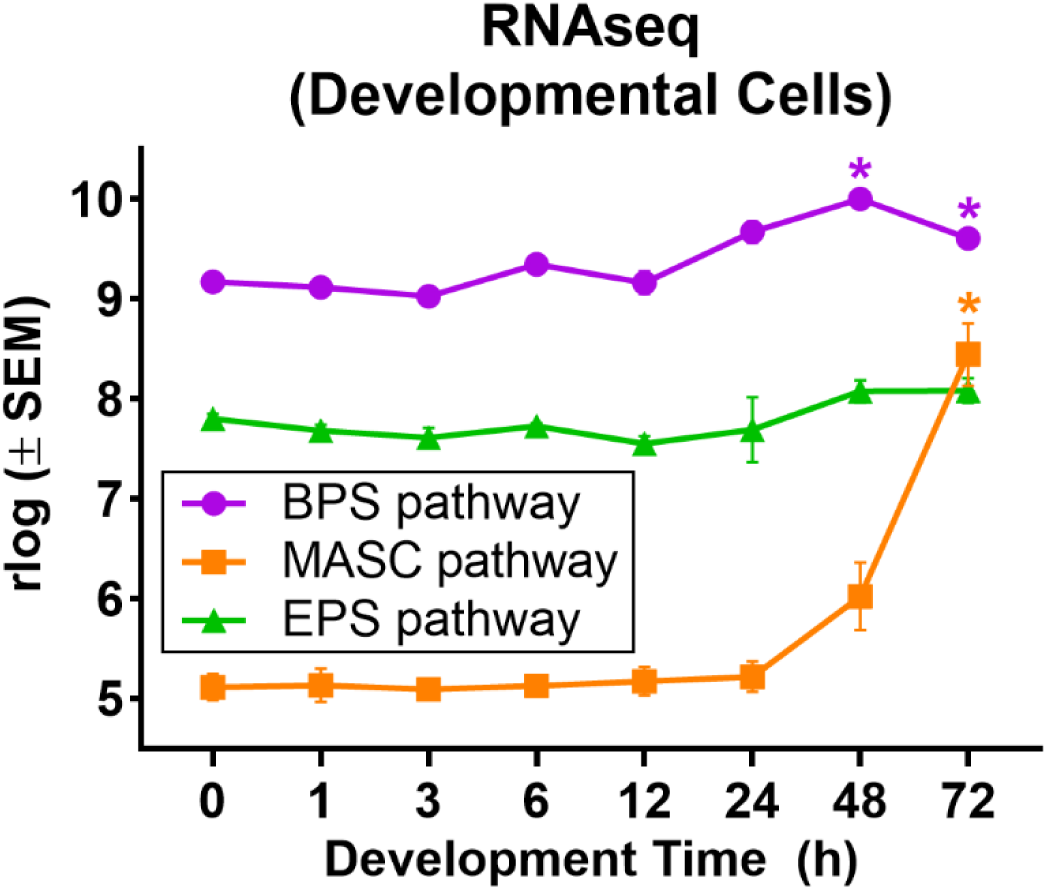
Expression levels of *M. xanthus* polysaccharide pathways during development. RNA was harvested from liquid broth-grown vegetative (t = 0 h) and starvation agar-grown developmental (t = 1 – 72 h) samples at various time points and subjected to RNAseq analysis^24^. Regularized log-transformed (rlog) expression values are displayed from triplicate measurements ± standard error of the mean (SEM) for expression levels of the Class-3 outer-membrane polysaccharide export (OPX) genes *wzaB*/*S*/*X* essential for BPS/MASC/EPS assembly-and-export^19,22,26^, respectively. The mean of each developmental time point was compared against the mean at 1 h via one-way ANOVA as this was the earliest on-agar reference point. Values with statistically significant differences (p ≤ 0.05) compared to 1 h are indicated (*).

### BPS-deficient myxospores are hyperaggregative

With the upregulation of the BPS pathway during sporulation periods of development **(Fig. 1)**, we next set out to probe the effect(s) of BPS on myxospore physiology. Natural myxospores were harvested from 5-day-old fruiting bodies produced by various polysaccharide-deficient mutant strains then imaged via phase-contrast microscopy. In all cases, myxospores were detected in a range of aggregate phenotypes, from isolated copies **(Fig. 2A)** to uncountable clumps **(Fig. 2B)**. However, the ratio of myxospores in smaller groups was heavily skewed toward uncountable aggregates for BPS^−^ myxospores that displayed considerable recalcitrance to disruption via sonication in a water bath **(Fig. 2B)**. Incidentally, exogenous treatment with a chemically-distinct biosurfactant (i.e. purified di-rhamnolipid-C_14_-C_14_ produced by *Burkholderia thailandensis* E264)^29^ resulted in partial disruption of the BPS^−^ myxospore aggregates **(Fig. 2B)**, confirming that the clumping phenotype characterized in these samples was indeed due to the biosurfactant activity of secreted BPS, analogous to the effect of BPS on vegetative cells^20^. Comparable results were obtained for chemically-induced myxospores (generated via glycerol treatment) **(Supplementary Fig. 2)**, pointing to an equivalent mode of aggregation between the two types of myxospores.

**Figure 2.**
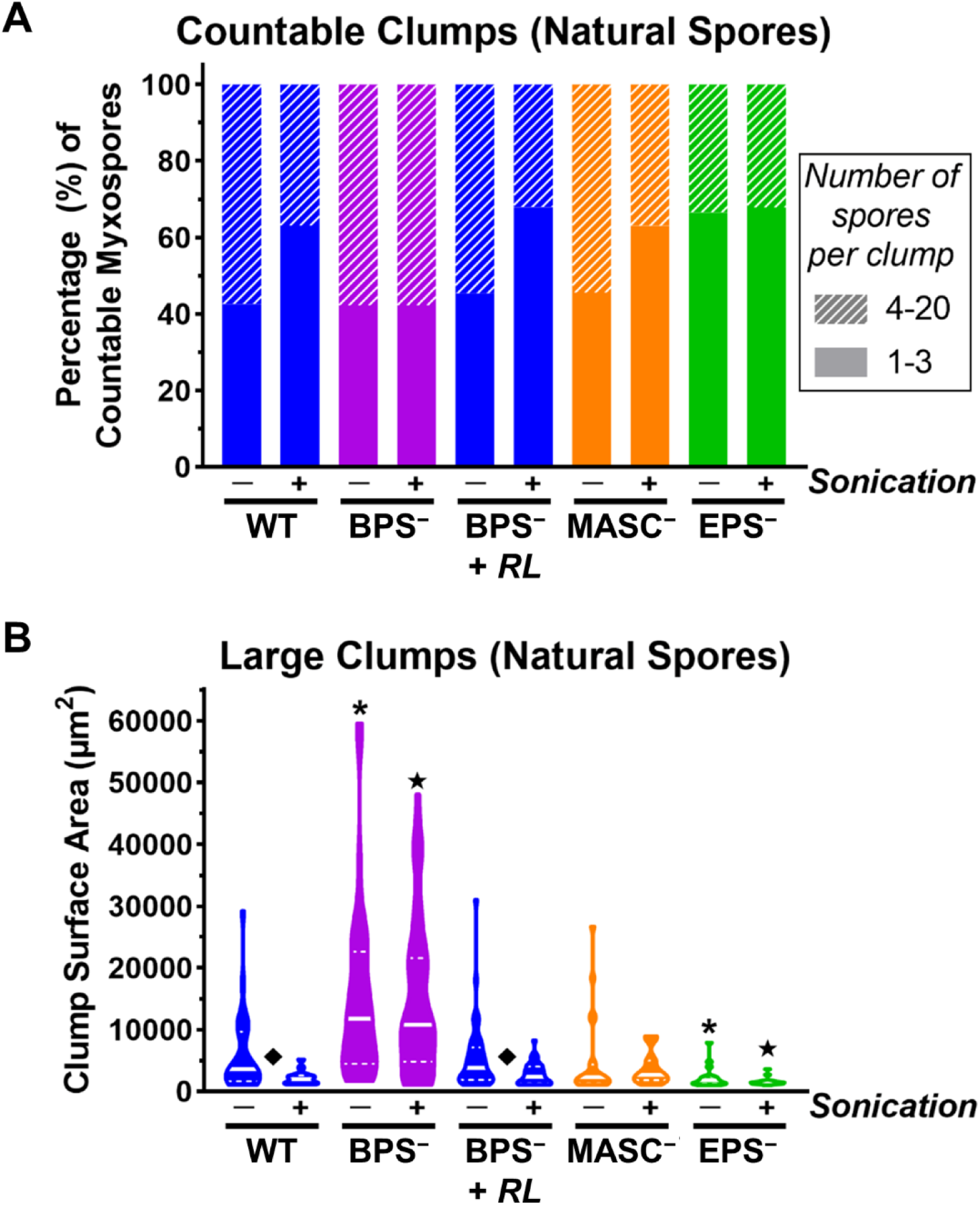
Aggregation profiles of natural myxospores. Samples were compared in the absence and presence of sonication in a benchtop waterbath for 30 s. BPS^−^ samples were screened in the absence and presence of rhamnolipid biosurfactant (+ *RL*). **A)** Percentage of countable myxospores, divided between groups of 1–3 and 4–20 myxospores per clump, as determined from phase-contrast microscopy imaging at 100×. **B)** Two-dimensional surface area of myxospore clumps ≥1000 µm^2^. Samples denoted with (i) an asterisk (**\***), (ii) star (⋆), and (iii) diamond (◆) displayed statistically significant (*p* ≤ 0.05) differences in distribution relative to (i) WT myxospores without sonication, (ii) WT myxospores with sonication, and (iii) each other, respectively, as determined via two-tailed Mann-Whitney U-tests.

### MASC-deficient myxospores display WT-like aggregation

Since the surface layer of myxospores has been attributed for over six decades to the presence of what is now termed MASC polysaccharide^14^, we initially hypothesized that secreted BPS was thus disrupting the outer-most layer on myxospores attributed to MASC polysaccharide (analogous to the BPS-induced destabilization of the EPS glycocalyx surrounding vegetative cells^20^). However, to our considerable surprise, both fruiting body-derived and glycerol-induced MASC^−^ myxospores displayed similar aggregative profiles pre-and post-sonication to those observed for WT myxospores **(Fig. 2, Supplementary Fig. 2)**, suggesting that MASC may not constitute the outer-most layer of either natural or induced myxospores after all.

### EPS-deficient myxospores are compromised for aggregation

In a previous investigation, Holkenbrink *et al.* (2014) reported that material with analogous chemical signatures to O antigen (typically capping LPS molecules) and/or secreted EPS co-purifies with spore-coat sacculi; however, these two signals were not distinguished from each other, and the detection was attributed to non-specific contamination with O antigen and/or EPS during the myxospore envelope isolation procedure^30^. Nonetheless, despite the extensive literature on myxospore formation and structure, we were unable to find past characterizations of myxospores from solely EPS-deficient strains. We thus included EPS^−^ myxospores in our aggregation analysis. Unexpectedly, myxospores derived from EPS^−^ cells displayed less clumping than both WT and MASC^−^ myxospores, with the difference even more pronounced compared to myxospores from BPS^−^ cells **(Fig. 2, Supplementary Fig. 2)**. These results suggest that EPS is present on the surface of myxospores and/or that secreted EPS is indeed acting like a non-specific glue to facilitate cohesion of myxospore aggregates.

### Cortex and/or coat layers are responsible for myxospore structural rigidity

To directly visualize myxospore surface morphology, we carried out transmission electron microscopy (TEM) imaging of natural myxospores. Fruiting bodies were scraped from starvation-agar plates and plunged directly into fixative solution to minimize any mechanical perturbation of myxospore surface architecture. Samples were then mechanically stabilized in agar, treated with osmium tetroxide to label lipid membranes, and treated with tannic acid to stabilize surface structures and enhance contrast of surface layers. As per standard TEM preparation steps, samples were dehydrated with ethanol, embedded in resin, sectioned, then subjected to heavy-metal staining (with uranyl acetate and lead citrate). Upon TEM imaging of WT myxospores, the IM, periplasmic, OM, and heavily-stained inner-and outer-cortex layers were all visible, as was an outer-most faintly-stained translucent layer surrounding the outer-cortex density^6–12^ **(Fig. 3)**. Importantly, no other defined layer was observed distal to this translucent surface layer. The thickness of this outer-most layer was relatively uniform around the WT myxospore perimeter, implying a non-random association between it and the myxospore surface. Moreover, this outer-most layer was found to make connections with the outer-most layers of nearby myxospores **(Fig. 3, *upper panel, asterisks*)**.

**Figure 3.**
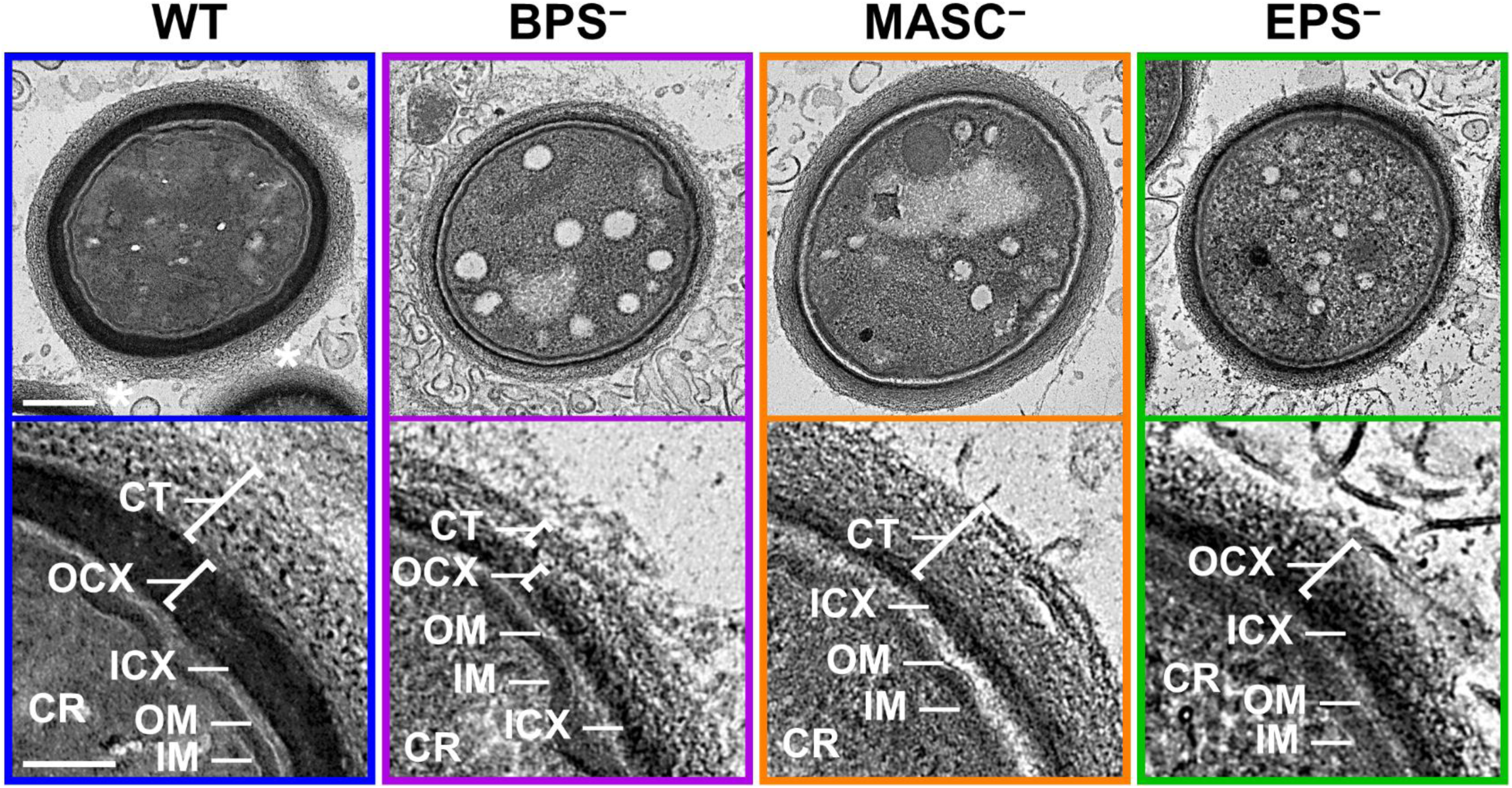
Transmission electron microscopy imaging of natural myxospore envelope architecture. Myxospores were obtained by scraping 5-day-old fruiting bodies and immediately immersing them in fixative solution. *Upper panels:* Whole-myxospore view. Scale bar = 300 µm. *Lower panels:* Magnified view of top-right quadrant from upper panels. Scale bar = 100 µm). CR, core; IM, inner membrane; OM, outer membrane; OCX, outer cortex; ICX, inner cortex; CT, coat.

In between the OM and outer-cortex layers of WT myxospores, the inner-cortex layer was visualized as a thin but darkly-stained ribbon of material. In instances when the IM and OM layers could both be seen to form invaginations after pulling away from the darkly-stained ribbon—likely due to dehydration from ethanol treatment during standard TEM sample preparation—this inner-cortex layer did not become invaginated and instead maintained its convex curvature, along with any outer-cortex and spore-coat material distal to it **(Fig. 3, *lower panel*)**. Since myxospores do not contain a rigid meshwork of peptidoglycan^31–33^, our data suggest that this darkly-stained inner-cortex ribbon may confer a degree of structural rigidity to the myxospore envelope and/or maintain its structure based on association with the layers (e.g. outer cortex and/or coat) distal to it. Interestingly, the inner cortex in natural myxospores could still be observed after only membrane labelling (via osmium tetroxide) and tannic acid treatment (i.e. in the absence of heavy-metal staining) **(Supplementary Fig. 3)**. Tannic acid binds strongly to proteins and acidic polysaccharides; it can act as a mordant, forming electron-dense complexes, and can thus create a strong contrast layer even before heavy-metal staining^34^. As such, this darkly-stained ribbon could be composed of acidic polysaccharide and/or proteinaceaous material. Importantly, unlike with the inner-cortex layer, no defined, contiguous tannic acid-stained layer was observed distal to the translucent myxospore coat layer, severely undercutting the notion of a potential protective protein coat surrounding the myxospore surface **(Fig. 3, Supplementary Fig. 3)**.

In glycerol-induced myxospores, the IM, periplasmic, and OM layers could still be reliably distinguished, as could the inner-cortex layer; however, in contrast to natural myxospores, there was a less clear stratigraphic distinction between darker-stained outer-cortex material and lighter-stained spore-coat material, with the two appearing to exist in a more mixed milieu **(Supplementary Fig. S4A)**. Finally, glycerol-induced myxospores were found to have a considerably thinner cortex and coat architecture compared to natural myxospores from fruiting bodies **(Fig. 3)**, consistent with previous reports^6,9,14^.

### BPS destabilizes the myxospore surface

In natural myxospores derived from BPS^−^ cells, the IM, periplasmic, and OM layers were visible, but with the dark outer-cortex and translucent surface-coat layers seemingly more compacted **(Fig. 3)**; this is similar to the more compact appearance of the glycocalyx layer in vegetative BPS^−^ *M. xanthus* cells^20^. Again, the darkly-stained inner-cortex ribbon distal to the OM layer in natural myxospores could still be seen following only osmium tetroxide and tannic acid treatment **(Supplementary Fig. 3)**. As BPS is the only net-negatively-charged *M. xanthus* polysaccharide identified to date^19^, the persistence of the darkly-stained inner-cortex layer even in the absence of BPS would support the contention that this ribbon is largely proteinaceous in nature. In glycerol-induced myxospores, the absence of BPS resulted in a splotchy appearance to the (sub)surface layers, with darker-staining likely outer-cortex material interspersed throughout the lighter-staining putative coat material. Despite their origin in shaking broth cultures, glycerol-induced myxospores generated in the absence of BPS were also still found to form close contacts with adjacent myxospores **(Supplementary Fig. 4A)**, consistent with the myxospore aggregation data discussed above. Taken together, these data point to BPS having a destabilizing effect on the myxospore surface analogous to its effects on the vegetative cell glycocalyx^20^.

### MASC polysaccharide constitutes the major sub-surface “cortex” layer of myxospores

Intriguingly, in natural MASC^−^ myxospores, the outer-most translucent layer was still visible while staining of the outer cortex layer was severely depleted **(Fig. 3)**. Moreover, using our minimally perturbative harvesting and preparation protocol, spherical myxospores were routinely obtained from our 5-day-old fruiting body preparations, even for MASC^−^ samples, countering previous reports of chemically-induced MASC^−^ myxospores being unable to stably form and instead reverting back to rod-shaped cells^30,35^. Though glycine amino acid side chains may be integrated into MASC polymers^9,30,31^, the darkly-stained inner-cortex ribbon above the OM layer was still present in the absence of MASC **(Fig. 3, Supplementary Fig. 3)**, suggesting that one or more as-yet-unidentified protein species may constitute the inner cortex. Nonetheless, our data indicate that MASC polysaccharide does not constitute the outer surface of myxospores, but rather the sub-surface outer-cortex layer, which sits atop a thin ribbon of what may be proteinaceous inner-cortex material.

### EPS constitutes the outer-most surface coat layer of myxospores

Contrary to MASC^−^ myxospores, EPS^−^ myxospores maintained heavy staining of the outer-cortex zone but no longer possessed the translucent layer corresponding to the outer-most surface of all other myxospores analyzed. This observation was consistent for both natural **(Fig. 3)** and glycerol-induced **(Supplementary Fig. 4)** myxospores. In addition, the surface of the outer-cortex layer appeared fuzzier and less dense in EPS^−^ myxospores compared to the interface between the outer-cortex and spore-coat layers observed in WT myxospores **(Fig. 3)**. As with all other strains tested, the darkly-stained inner-cortex ribbon distal to the OM layer in natural myxospores was still present in the absence of EPS **(Fig. 3, Supplementary Fig. 3, Supplementary Fig. 4)**. Crucially, these findings provide convincing evidence that the outer-most layer of myxospores may in fact be composed of the same EPS that constitutes the vegetative cell glycocalyx. Moreover, the even and well-defined structuration of this outer-most layer argues against the notion that EPS on the myxospore surface is simply a non-specific amorphous contaminant, and instead promotes the surface presence of EPS as an intentional outcome of myxospore biogenesis. Finally, our data are consistent with the outer-cortex layer being secreted under an existing EPS layer, resulting in compaction of the former under the latter.

### EPS constituting the myxospore coat is analogous to that surrounding vegetative M. xanthus cells

Numerous techniques to analyze the nature of cell-surface EPS in *M. xanthus* have been implemented, all of which are dependent on well-established changes in cell-surface properties. As such, we first adapted the physicochemical MATH (microbial adhesion to hydrocarbons) test^20,36^ for use with myxospores from WT and the various polysaccharide-deficient mutant strains to characterize differences in relative surface hydrophobicity. In this test, spore (or cell) samples in aqueous medium are mixed with hexadecane, and hydrophobic interactions are inferred from the reduction in OD_600_ of the aqueous phase following phase separation of the emulsion^37^. The degree of partitioning between samples is directly impacted by the overall amount, accessibility, and composition of surface material^20^. Compared to WT samples, myxospores from BPS^−^ cells partitioned less efficiently with the hexadecane, similar to vegetative BPS^−^ cells that have the same amount of surface glycocalyx material as WT, but for which the glycocalyx is more compacted and less accessible **(Fig. 4A)**. Intriguingly, natural MASC^−^ myxospores displayed no significant differences in relative hydrophobicity compared to WT myxospores **(Fig. 4A)**, pointing to analogous properties of their respective outer-most layers; however, this equality was lost in glycerol-induced myxospores **(Supplementary Fig. 5A)**, potentially reflecting overall differences in MASC vs EPS distribution in the envelope of induced myxospores **(Supplementary Fig. 5A)**. Incidentally, fewer EPS^−^ myxospores partitioned with the hexadecane **(Fig. 4A, Supplementary Fig. 5A)**, consistent with results from vegetative EPS^−^ cells^20^, indicating a depleted surface polysaccharide layer^20^.

**Figure 4.**
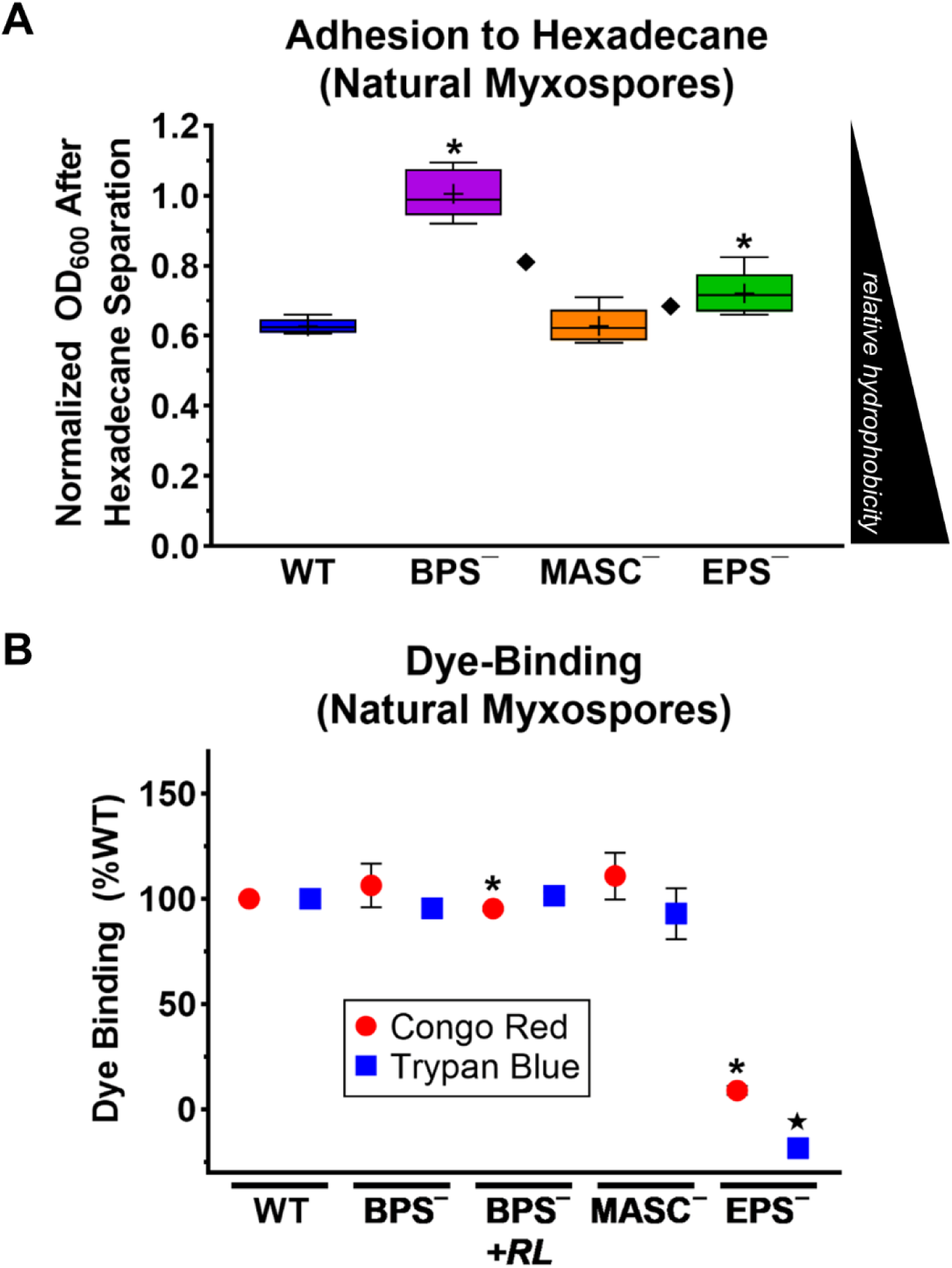
Surface properties of natural myxospores. **A)** OD_600_ readings of cuvettes containing myxospores in aqueous buffer mixed with hexadecane, after 1 h of emulsion separation. Lower and upper box limits denote 25^th^ and 75^th^ percentile values, with the lower/upper Tukey fences, median (solid line), and mean (+) values indicated. Samples denoted with (i) an asterisk (**\***) and (ii) a diamond (◆) displayed statistically significant differences in distribution relative to (i) WT myxospores (*p* ≤ 0.0159) and (ii) each other (*p* ≤ 0.0317), respectively, as determined via two-tailed Mann-Whitney U-tests (n = 5). **B)** Mean values (n = 3) for Congo Red and Trypan Blue dye binding relative to WT myxospores, displayed ± standard error of the mean. Mutant myxospore samples denoted with (i) an asterisk (**\***) and (ii) star (⋆) displayed statistically significant differences in mean values compared to the WT Congo Red (*p* ≤ 0.0131) and Trypan Blue (*p* < 0.0001) samples, as determined via two-tailed Student’s t-tests. *RL*: Samples treated with rhamnolipid biosurfactant prior to dye treatment.

We next carried out binding comparisons using the aromatic diazo dyes Congo Red and Trypan Blue, both extensively used for *M. xanthus* EPS detection^38–40^. Using WT myxospores as a reference, MASC^−^ myxospores displayed no significant differences in the levels of bound dye, with similar results obtained for BPS^−^ myxospores. Analogous to vegetative cells^19,39^, for EPS^−^ myxospores, binding of either dye was abolished **(Fig. 4B, Supplementary Fig. 5B)**, indicating a lack of suitable glycocalyx material on the surface of these myxospores.

Finally, to demonstrate the functionality of EPS on myxospores, we creatively exploited its status as a “shared good” within *M. xanthus* communities^19,23^. Swarm spreading via T4P extension and retraction^41^ is dependent on the presence of EPS on adjacent cells, as this EPS interacts with the T4P^42^ and acts as a retraction signal^43^. While EPS^−^ swarms are intrinsically impaired for T4P-dependent spreading, supplementation of these swarms—with either isolated EPS glycocalyx material^23^ or EPS-producing cells^19^—transiently restores EPS-dependent spreading of the entire swarm via T4P function. We first engineered EPS^−^ cells to express secreted red-fluorescent mCherry fused to an IM signal sequence, which results in mCherry localized to the periplasm, but tethered to the periplasmic leaflet of the IM (IMss-mCherry)^44^. Using EPS^−^ cells expressing IMss-mCherry, we added myxospores from our panel of WT and polysaccharide-deficient strains, and probed for re-activated T4P-dependent swarm spreading of red-fluorescent EPS-deficient cells on soft (0.5% w/v) CYE agar. Importantly, imaging of spreading re-activation was limited to 6 h post-inoculation of the agar medium with the cell–spore mixtures; this is a time at which germinating myxospores are undergoing conversion to rod-shaped cells but no RNA synthesis has taken place indicating a lack of metabolic activity^45^. This ensured that EPS-elaborating spores did not germinate, begin to replicate, produce new EPS, and undertake T4P-mediated spreading of their own. Also, as *M. xanthus* is known to physically exchange OM lipids and proteins (via OM–OM fusion events between compatible cells)^3^, our use of mCherry targeted to the IM (instead of to the OM or soluble in the periplasm) ensured that no fluorescent signal could be transferred to cells that had newly-germinated from the introduced myxospores.

Under these conditions, myxospores from all strains tested, including those that were MASC^−^, were able to induce formation of cohesive swarm fronts by intrinsically EPS^−^ red-fluorescent cells and re-activate EPS-dependent collective cell migration outward from these swarm fronts. The only exception was EPS^−^ myxospores, the addition of which had no effect on swarm-edge cohesion or re-activation of collective motility **(Fig. 5)**. Of note, re-activated swarm-front cohesion and outward motility were not equivalent among the different introduced myxospore types, suggesting that EPS on the surface of WT, BPS^−^, and MASC^−^ myxospores is differently held by their respective spore surfaces.

**Figure 5.**
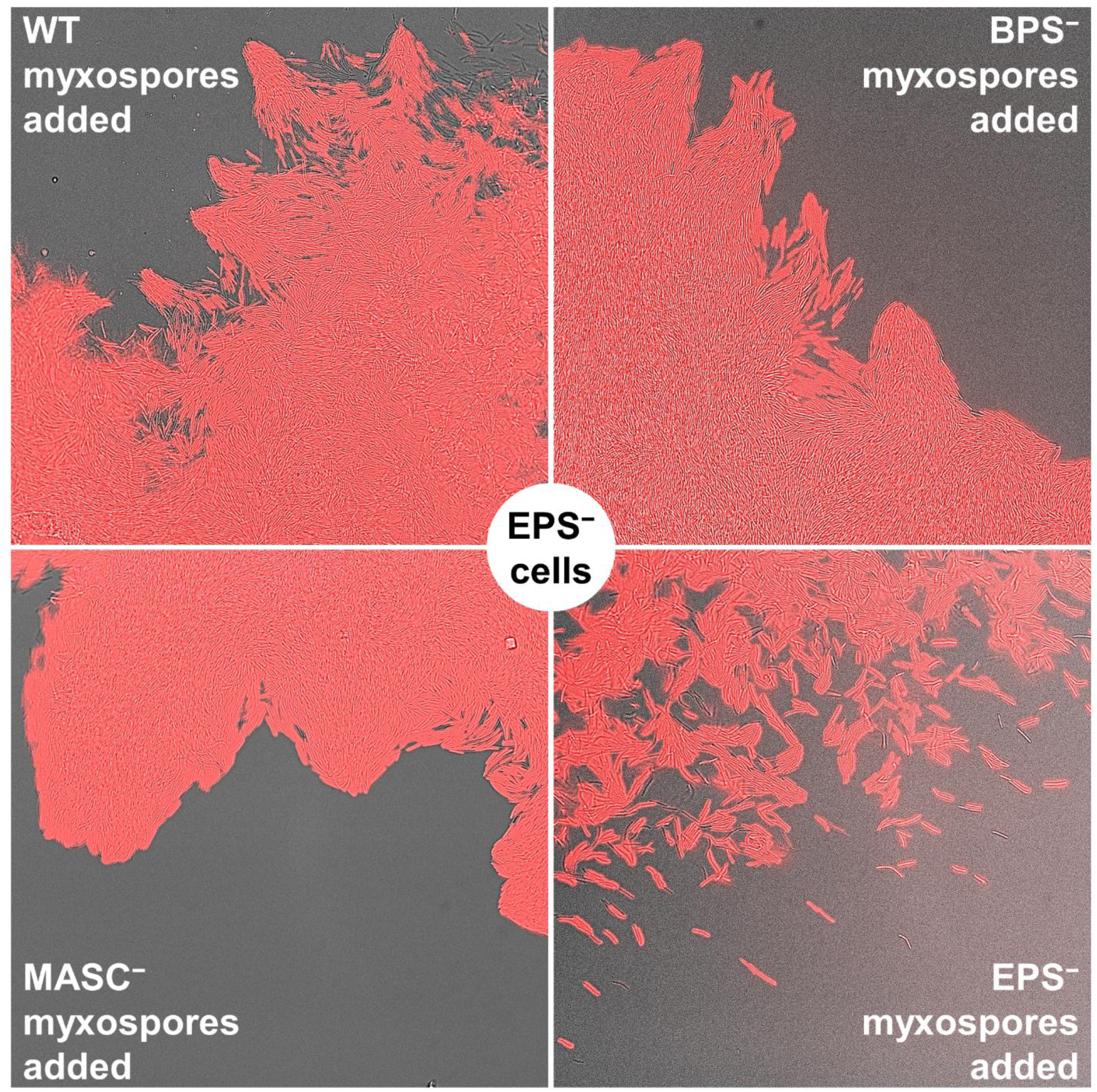
Reactivation of EPS-dependent T4P-mediated group motility in EPS^−^ cells via mixing with EPS^+^ myxospores. EPS-deficient cells of *M. xanthus* Δ*wzaX*, expressing IMss-mCherry, were mixed with myxospores derived from different polysaccharide-mutant *M. xanthus* strains. Cells and myxospores were spotted together on “soft” 0.5% (w/v) CYE agar to favor T4P-dependent swarm spreading, and were imaged after 6 h at 32 °C under a 40× objective. Displayed is an overlay between the phase-contrast and red-fluorescence images taken at the swarm edge. Mixtures were assayed for the appearance (or lack thereof) of contiguous swarm fronts and/or the formation of leading edge rafts/projections of cells, all indicative of tight T4P-dependent inter-cell interactions and group motility that depend on the presence of EPS on the cell surface.

Ultimately, the surface hydrophobicity, dye-binding, and strain supplementation results collectively confirm that, instead of MASC polysaccharide, the outer-most layer of myxospores is indeed composed of the same EPS as that on the surface of vegetative cells. This is consistent with the previous co-purification of EPS material during isolation of spore coat material from chemically-induced myxospores^30^. As such, we also propose that the term “MASC” should now instead denote the “<u>ma</u>jor <u>s</u>pore <u>c</u>ortex” polysaccharide **(Fig. 6)**.

**Figure 6.**
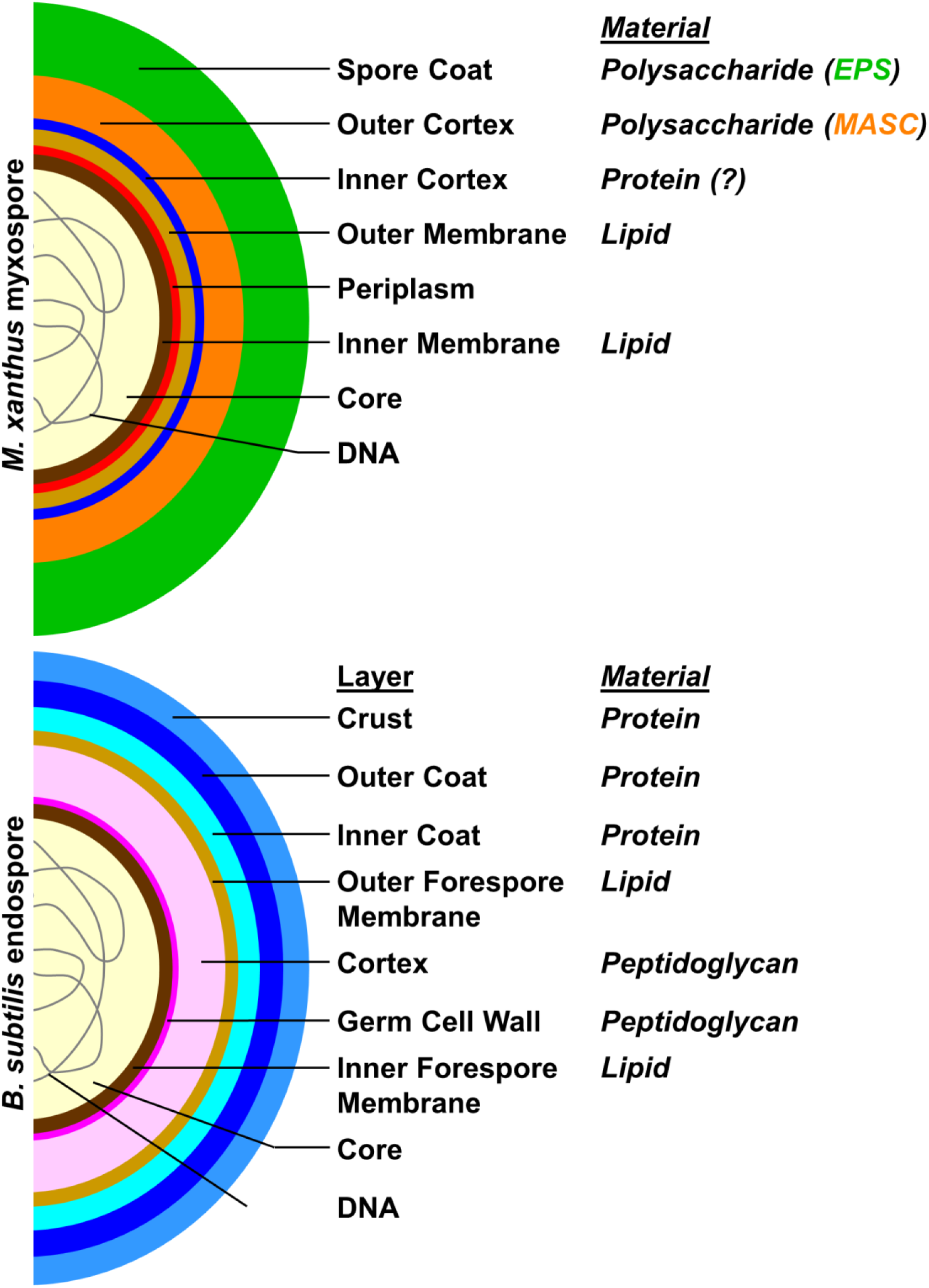
Revised bacterial spore envelope architecture. Based on the findings herein, the myxospore outer-coat is composed of EPS, while the MASC polysaccharide forms a sub-surface outer-cortex layer. This sits atop an inner-cortex layer that may be composed of one or more protein species. The architecture of *Bacillus subtilis* endospores (composed of lipid, peptidoglycan, and protein) is depicted for comparison.

## DISCUSSION

In this study, we examined the surface properties and aggregation behavior of *M. xanthus* myxospores, with a particular focus on the contributions of EPS, BPS, and MASC polysaccharide to these processes. Our findings challenge long-standing assumptions about the architecture of the myxospore surface and highlight the nuanced interplay between polysaccharide composition and biosurfactant activity in regulating spore clustering and dispersal.

We found that myxospores formed from BPS-deficient cells are hyperaggregative, forming persistent clumps that resist physical disruption. As with vegetative cells, and the complementation *in trans* using exogenous biosurfactant, our findings reinforce the notion that BPS acts to modulate the presentation and/or accessibility of surface polysaccharides. From an ecological and developmental perspective, the regulation of myxospore aggregation by BPS has important implications. While aggregation within fruiting bodies likely confers protection and may coordinate germination, excessively tight packing of spores could limit the diffusion of nutrients or signaling molecules necessary for synchronized reactivation. By functionally destabilizing EPS and reducing surface stickiness, BPS may help maintain optimal spacing between spores (similar to its effects on vegetative cells^20^, facilitating more uniform exposure to environmental cues and promoting collective emergence of vegetative cells. This insight suggests that BPS-mediated modulation of EPS is not merely a matter of individual spore dispersal but represents a key factor in the spatial organization and functional success of multicellular structures.

Our analysis also clarifies the long-standing question of myxospore surface composition. Contrary to the prevailing view until now that MASC polysaccharide forms the outermost layer and that EPS is a non-specific contaminant, our imaging, hydrophobicity, dye-binding, and spreading-complementation assays indicate that MASC instead constitutes the sub-surface outer-cortex layer, while EPS forms the true external surface. This external surface was previously shown to form connections between adjacent myxospores^6^, an observation we have confirmed in this investigation, suggesting a possible structural role for this myxospore layer in the final architecture of macroscopic fruiting body assemblies.

A logical explanation for the presence of EPS on mature myxospores is that these spores derive directly from vegetative cells that already possess EPS on their surfaces, and that the differentiation process does not remove this pre-existing layer. Thus, the bulk of EPS is likely not newly synthesized during sporulation but rather inherited, consistent with our observations that EPS^−^ spores completely lack the surface coat layer and fail to complement EPS-dependent swarm functions. As developing myxospores already start out surrounded by an EPS “shield”, subsequent secretion of MASC polysaccharide would result in accumulation of this myxospore-specific polymer under the existing EPS layer, explaining the presence of the sub-surface presence of the outer (major) cortex layer in myxospores imaged by ourselves and others^6,7,12,46^.

The presence of three clearly-defined layers distal to that of the OM in myxospores was only ever reported previously by Inouye *et al* (1979), without knowing the identities of the various constituents. Therein, the outer-most layer (now shown to be EPS) was reported to be removable following treatment with 1M NaCl^6^, suggesting that the EPS layer is not as tightly held by the spore as the underlying MASC layer. Incidentally, the MASC polysaccharide layer on myxospores is known to exist in a compacted state. As discussed above, part of this compaction may occur from MASC being secreted under an existing polysaccharide layer. However, it has been shown that following MASC polysaccharide secretion, the polymer becomes wound around the myxospore akin to loose yarn being wound tightly around a ball. This process is mediated by proton motor-powered migration of the trans-envelope Nfs complex around the spore periphery^47^, a system with direct functional paralogies with the Glt gliding motility apparatus^17,18^ that powers *M. xanthus* single-cell locomotion. As such, the relatively even distribution of EPS atop the MASC polysaccharide outer-cortex layer could be an indirect result of the subsurface MASC polymers being tightened, and the EPS polymers, in essence, being dragged along for the ride. In the future, a comparative analysis of Nfs function between WT and BPS^−^ myxospores would provide intriguing insights into the role of spore-coat rigidity in the efficient distribution of MASC around the myxospore subsurface.

Density corresponding to the inner cortex described herein was also reported by Inouye *et al* (1979), with our differential staining and TEM imaging suggesting this layer may be composed of one or more protein species. Several proteins associated with the myxospore envelope have been previously reported in the literature. Protein S has long been associated with the surface of myxospores from fruiting bodies, but not those produced via glycerol induction^6,48^. Moreover, it is not required for the biogenesis or viability of myxospores, it can be gently washed from the myxospore surface, and it may instead function within fruiting bodies to influence myxospore adhesiveness^49^. Protein C has similarly been deemed a surface protein in fruiting-body myxospores (but not for glycerol-induced myxospores); however it too could be washed from the myxospore surface^50^ and later was identified to be simply an accumulated proteolyzed fragment of the FibA zinc metalloprotease that associates with the EPS material of vegetative cells^13,51^. Finally, Protein U, which is a secreted protein^52^ detected in the extracellular matrix already at 24 h during development^53^, can also be removed from the myxospore surface. Moreover, it is not required for reconstitution of the cortex zone in myxospores that have had their surface layers removed then reconstituted through *in trans* addition of purified myxospore surface material^6^. Proteins S, C, and U are thus highly unlikely to be structurally-important inner-cortex proteins and instead appear to non-specifically associate with the now-identified EPS surface coat layer on myxospores during development in fruiting body structures.

Conversely, CbgA is a candidate for an inner-cortex protein. It was initially identified via its sequence homology to SpoVR, a key protein involved in endospore cortex formation in *B. subtilis*^54^. Cells deficient in CbgA generated myxospores with either thin cortex layers (i.e. possibly lacking the MASC outer cortex), or which lacked them outright (with no distinction made therein between inner-vs. outer-cortex layers), while still maintaining the same levels of coat material (now known to be EPS). These CbgA-lacking myxospores displayed greater temperature-and detergent-sensitivity than WT myxospores^12^. Through proteomic analysis of fruiting body myxospores, three additional candidates were identified: MspA, MspB, and MspC (the coding sequences for which are separated by >160 and >4537 genes, respectively). Mutant strains knocked out for the respective genes were still able to aggregate and produce myxospores in fruiting bodies that elaborated myxospore coat (i.e. EPS) layers. However, cortex formation was severely impacted in all three mutants. Strains deficient in either MspA or MspB generated myxospores that produced staining comparable to our MASC^−^ mutant (i.e. with a heavily-stained intact inner-cortex layer, but lacking the outer-cortex layer under the coat). Cells lacking MspC produced myxospores more consistent with an absent inner-cortex layer (i.e. no heavy OM-adjacent staining, but still with a dark, slightly more diffuse, outer-cortex layer under the coat). Mutants in all three genes displayed increased sensitivity to heat and detergent, while MspC-deficient myxospores were also more sensitive to sonication^7^. Ultimately, one or more of these four proteins could be localized to the cortex zone of myxospores.

A notable outcome of this work is the sharp contrast between the revised architecture of *M. xanthus* myxospores and the canonical architecture of *B. subtilis* endospores **(Fig. 6)**. In *B. subtilis*, the spore surface is defined by a multilayered proteinaceous coat and, in some cases, a glycoprotein-rich exosporium, structures synthesized *de novo* during sporulation. In contrast, our data indicate that in *M. xanthus*, the outermost surface is likely not newly elaborated but instead derived from pre-existing vegetative EPS that is retained through the differentiation process. This fundamental distinction highlights divergent strategies for achieving surface protection: while *B. subtilis* invests in constructing rigid, protein-based barriers, *M. xanthus* relies on polysaccharides that are dynamically modulated by biosurfactant activity. The result is a more “inherited” and chemically flexible surface, whose properties can be tuned by extracellular factors such as BPS. These differences underscore the evolutionary diversity of spore-protection strategies across bacterial lineages and suggest that diderm developmental systems may have prioritized adaptability and social regulation over the rigid, individually focused defense mechanisms typified by endospores.

Overall, our results redefine the surface architecture of *M. xanthus* myxospores and provide a mechanistic explanation for how biosurfactant-mediated EPS remodeling influences both single-spore and population-level behaviors. By revealing that EPS (rather than MASC) constitutes the outermost surface layer, and by demonstrating the common effects of BPS on vegetative cells and dormant spores, we uncover fundamental principles governing spore dispersal, aggregation, and readiness for germination. These findings open the door to further studies on how polysaccharide composition and surface-active molecules coordinate multicellular development in diderm bacteria.

## MATERIALS & METHODS

### Growth of bacterial cells

Myxospores from frozen stocks were germinated on CYE medium (1% Bacto Casitone [w/v], 0.5% yeast extract [w/v], 0.1% MgCl₂, 10 mM MOPS, pH 7.4) solidified with 1.5% (w/v) Bacto Agar, incubated at 32 °C. Vegetative cultures were initially inoculated from these plates into 12.5 mL CYE broth in 125-mL flasks and grown overnight at 32 °C with shaking (220 rpm).

### Spore generation and harvesting

To obtain natural spores, the optical density read at 600 nm (OD_600_) was first measured via spectrophotometer for vegetative cell cultures transferred to a polystyrene cuvette. Cells were then sedimented via centrifuge (16 000 × *g*, 5 min) and resuspended to an OD_600_ of 5.0 in TPM buffer (10 mM Tris-HCl [pH 7.6], 1 mM KH₂PO₄, 8 mM MgSO₄). This resuspension was spotted (5 µL) on CF-agar (0.015 g Bacto Casitone peptone, 4 mL MOPS [1 M, pH 7.6], 0.4 mL K₂HPO₄ [1 M, pH 7.6], 4 mL MgSO₄·7H₂O [0.8 M] starvation plates, solidified with 1.5% [w/v] BactoAgar), and grown at 32 °C for 120 h (i.e. 5 days) to promote fruiting body formation. Using a sterile inoculating loop, nine to twelve separate fruiting bodies from these plates were scraped and pooled into 1 mL TPM buffer, followed by OD_600_ analysis, centrifugation (see above), and resuspension of the myxospores in TPM buffer to the desired OD_600_ depending on the assay.

For chemically-induced spores, *M. xanthus* cells were first grown overnight in CYE broth under vegetative culture conditions (see above), then used to inoculate 3 mL CYE cultures in sterile glass tubes to an initial OD_600_ of 0.25. To induce myxospore formation, 109.4 µL of 100% glycerol was added to each subculture^55^, followed by incubation at 32 °C with shaking (220 rpm) for an additional 24 h. The OD_600_ for these glycerol-treated subcultures was then determined, followed by centrifugation (see above), and resuspension of the chemically-induced myxospores in TPM buffer to the desired final OD_600_ based on the specific assay.

### Spore clump analysis

Natural and chemically-induced myxospore samples resuspended in TPM buffer at an OD_600_ of 5.0 were spotted (0.75 µL) on untreated long glass coverslips, overlaid with a TPM 1.5% agar (w/v) pad, then imaged at 40× and 100× magnification under phase-contrast illumination (20% LED illumination intensity, 10-ms exposure time, binning mode: 2 × 2) on a Zeiss AxioObserver 7 fluorescence microscope. To test clump stability, myxospore resuspensions in 1.5 mL microfuge tubes were placed in a Branson ultrasonic water bath for 30 s prior to spotting on glass slides. Myxospores were counted individually for clumps containing 20 or fewer visible myxospores. For larger aggregates, 2-dimensional aggregate surface area was measured using the MicrobeJ module^56^ for Fiji image analysis software.

### Transmission electron microscopy

For TEM imaging of natural myxospores, 5-day-old fruiting bodies (see above) were scraped directly into 1 mL fixative solution (2% paraformaldehyde and 2% glutaraldehyde, in 0.1 M cacodylate buffer containing 0.2 M sucrose) and incubated without shaking for 12 h at 4 °C. Fruiting bodies were then gently sedimented (1 000 × *g*, 5 min) and resuspended three times in 0.1 M cacodylate buffer containing 0.2 M sucrose before being post-fixed in 2% OsO_4_ in 0.1 M cacodylate buffer for 2 h at room temperature. Next, samples were sedimented and resuspended three times in 1 mL ddH_2_O for wash steps, then resuspended in 1% tannic acid solution for 1 h at room temperature. This procedure helped preserve the natural surface architecture of any myxospores present. Samples were then dehydrated with a series of 10-min steps using 30, 50, 70, and 95% ethanol before the final three 10-min dehydration steps using 100% ethanol. Subsequently, samples were infiltrated in a stepwise fashion as follows: ethanol-Spurr resin (1:1) for 1 h at room temperature, 100% Spurr resin for 2 h at room temperature, then fresh 100% Spurr resin overnight at room temperature. Samples were then embedded in Spurr resin before incubation at 60–65 °C for 20–30 h. After polymerization, samples were sectioned (60–90 nm) using an ultramicrotome (Leica EM). Sections were collected on copper 200-mesh grids then stained with 5% uranyl acetate in 50% ethanol (15 min) followed by Reynold’s lead citrate (5 min). Imaging was carried out using a Hitachi HT7800 transmission electron microscope with AMT NanoSprint15 Mk-II camera. For glycerol-induced myxospores, samples were harvested and processed as detailed above.

### Cell-surface hydrophobicity testing

MATH testing of myxospores was adapted from a previous protocol used for vegetative cells^20^. In brief, *M. xanthus* myxospores (natural or induced) were prepared as described above, sedimented via centrifuge (16 000 × *g*, 5 min), and resuspended via aspiration with a p1000 micropipette in 1 mL CYE broth to a calculated OD_600_ of 1.0 in a 2-mL microfuge tube. This volume was then transferred to a spectrophotometer cuvette, after which the OD_600_ was determined for the spore resuspension (termed “Mix 1”). Cuvette contents were subsequently transferred back to the microfuge tube, mixed using a vortex mixer (20 s), then transferred back to the cuvette after which OD_600_ was read again (termed “Mix 2”). This step served as an internal control to show that any subsequent changes in OD600 reading were not simply due to further spore dispersal due to vortex mixing. Cuvette contents were again transferred back to the microfuge tube, after which 75 µL of hexadecane (Sigma) was added to the spore resuspension, followed by vortex mixing (20 s). This emulsion was then transferred back to the spectrophotometer cuvette (termed “Mix 3”). Cuvettes were sealed with Parafilm and left in a cuvette rack on the benchtop without disruption for 1 h to allow for emulsion separation. After the 1 h period, the OD600 of samples in the cuvettes was re-determined (termed “Mix 4”). All “Mix 4” values across all strains and replicates were first normalized to their respective “Mix 1” values, followed by a second normalization of the different mutants’ values relative to the wild-type control for a given replicate.

### Congo Red and Trypan Blue dye retention

Congo Red-binding and Trypan Blue-binding analyses were carried out according to a protocol modified from that used with vegetative cells^19^. Natural and glycerol-induced myxospores were obtained as described above, sedimented via centrifuge (16 000 × *g*, 5 min), and resuspended in 1 mL of TPM buffer to an OD_600_ of 1.0. A 900-µL aliquot of this resuspension was mixed with 100 µL of stock Congo Red (150 µg/mL) or stock Trypan Blue (100 µg/mL) solution in 2-mL microfuge tubes. Sample tubes were laid on their sides, covered with aluminum foil to protect from light, incubated for 1 h at room temperature on a nutator platform, then sedimented (16 000 × *g*, 5 min). From the clarified supernatant, 900 µL were transferred to a polystyrene cuvette. Using myxospore-free “TPM + dye” (i.e. 900 µL TPM + 100 µL Congo Red/Trypan Blue stock solution) control samples, a spectrophotometer was then blanked at an absorbance of 490 nm (A_490_) for Congo Red, and 585 nm (A_585_) for Trypan Blue. The A_490_ and A_585_ were subsequently read for the Congo Red-and Trypan Blue-containing clarified supernatants, respectively. Absorbance values were then normalized to the A_490_ and A_585_ values for the WT sample for each biological replicate experiment.

### Spore mixing to transiently complement EPS-deficient cells

The day prior to mixing, a square (12 cm × 12 cm × 1.7 cm) plastic Petri dish was filled with 50 mL CYE 0.5% (w/v) “soft” agar, left uncovered in the biohood to cool for 15 min, re-covered, then stored face-down on the benchtop at room temperature overnight. The day of the experiment, vegetative overnight cultures of EPS-deficient *M. xanthus* expressing red-fluorescent IMss-mCherry^44^ **(Supplementary Table S1)** were resuspended in TPM buffer to an OD_600_ of 5.0. Natural and chemically-induced myxospore samples from non-fluorescent strains were also resuspended in TPM buffer to an OD_600_ of 5.0. In a biohood, on the previously-poured plate, 5 µL of IMss-mCherry-expressing EPS-deficient cells were spotted and left uncovered for 10 min to dry. On the same spot, 5 µL of non-fluorescent natural/induced myxospores were added and allowed to dry for another 10 min. Plates were then sealed with Parafilm and incubated at 32 °C. After 6 h of mixing, using a sterile scalpel, a square of agar under the mixed spots was excised and placed face down on an untreated long glass microscope cover slip. For detection of red-fluorescent mCherry signal, slides were imaged at 40× magnification with an alpha Plan-Apochromat oil-immersion objective (captured with 50% LED illumination intensity, 150-ms exposure time, binning mode: 2 × 2, reflector: 45 Texas Red) on a Zeiss AxioObserver 7 inverted fluorescence microscope with an Axiocam 512 camera. Phase contrast imaging was carried out with the same objective (20% LED illumination intensity, 10-ms exposure time, binning mode: 2 × 2). Images from the different channels were overlaid in Fiji, with image panels assembled in Powerpoint (sharpened +50%).

### RNA-seq analysis of OPX polysaccharide export gene expression

To assess transcriptional changes during vegetative growth and multicellular development in *Myxococcus xanthus*, we re-analyzed published RNA-seq data from Sharma *et al.* (2021) following the complete developmental time course^24^. Expression values were extracted for vegetative and defined developmental time points up to 72 h, with a specific focus on the transcription of the Class-3 outer-membrane polysaccharide export (OPX) polysaccharide biosynthesis genes *wzaB* (BPS pathway; MXAN_1915; MXAN_RS09285), *wzaS* (MASC pathway; MXAN_3225; MXAN_RS15620), and *wzaX* (EPS pathway; MXAN_7417; MXAN_RS35880). In the original study^24^, raw FASTQ files were quality-checked using FastQC^57^ trimmed with Trimmomatic^58^, aligned to the *M. xanthus* genome using Bowtie^59^, with gene counts generated via HTSeq-count^60^. Differential expression analysis was performed using DESeq2^61^, with regularized log-transformed (rlog) expression values were obtained using the rlog() function in DESeq2. For statistical analyses, one-way ANOVA analysis was performed, with each row representing matched data, assuming Gaussian distribution of residuals, with Geisser-Greenhouse correction applied (i.e. not assuming sphericity). The mean of each column was thusly compared against the mean of a control column (which was set to the 1 h time point as that is the earliest time point for samples harvested from agar).

