## Supplementary Figures for "Biosurfactant polysaccharide drives dispersal of exopolysaccharide-coated myxospores in *Myxococcus xanthus*"

*corresponding author

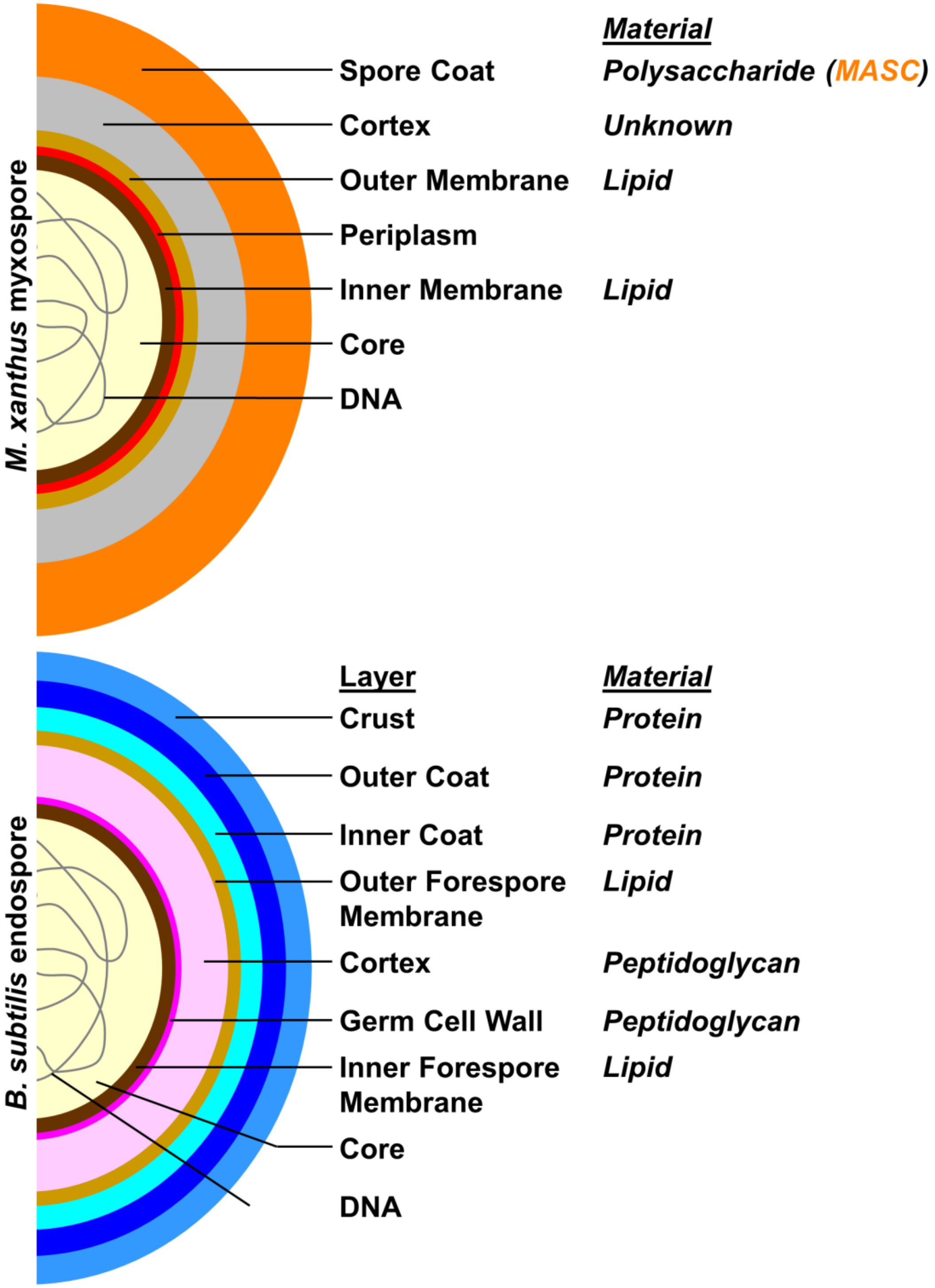

**Supplementary Figure 1. Current models of bacterial spore envelope architecture.** Previously-proposed *Myxococcus xanthus* (diderm bacterium, i.e. "Gram negative") myxospore layers (composed of lipid and polysaccharide), compared to known *Bacillus subtilis* (monoderm bacterium, i.e. "Gram positive") endospore layers (composed of lipid, peptidoglycan, and protein).

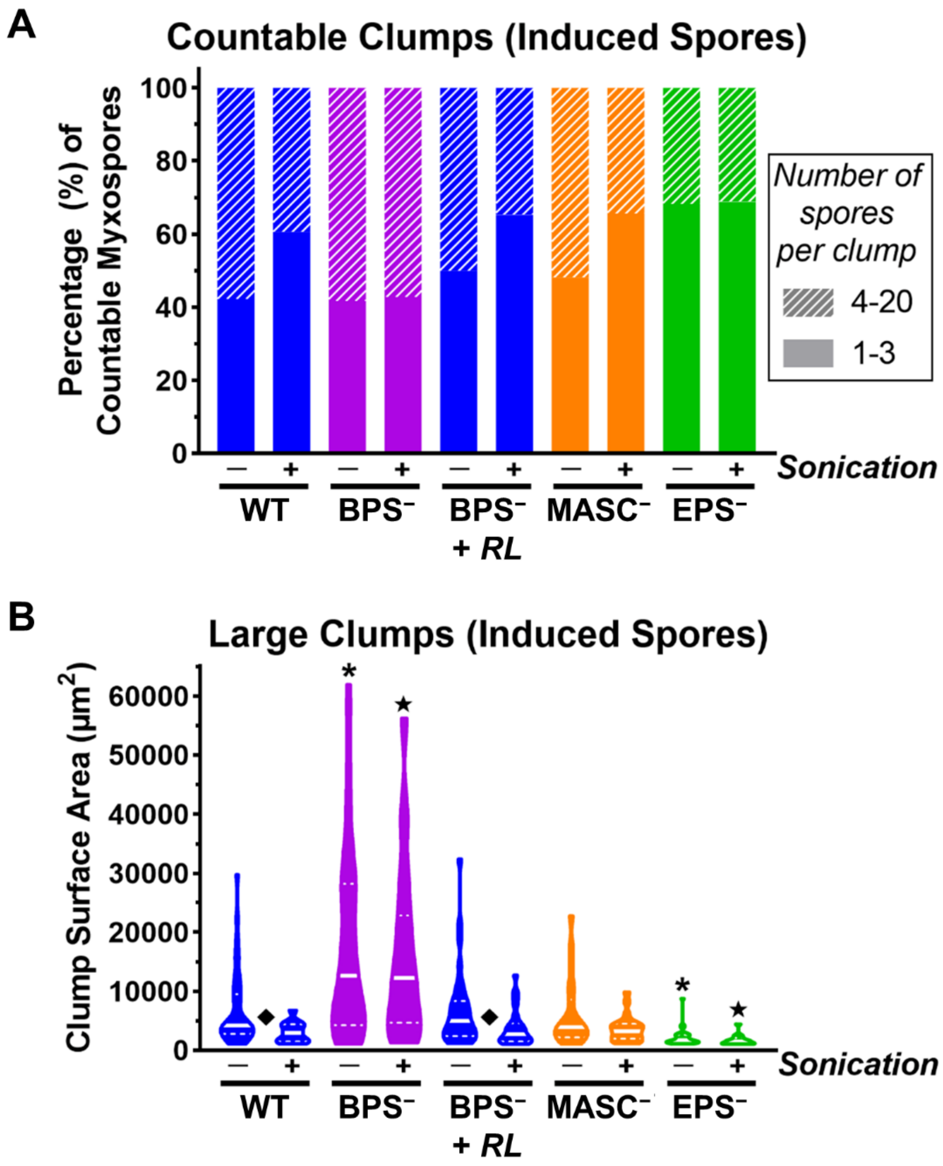

**Supplementary Figure 2. Aggregation profiles of glycerol-induced myxospores.** Samples were compared in the absence and presence of sonication in a benchtop waterbath for 30 s. BPS⁻ samples were screened in the absence and presence of rhamnolipid biosurfactant (+ *RL*). **A)** Percentage of countable myxospores, divided between groups of 1–3 and 4–20 myxospores per clump, as determined from phase-contrast microscopy imaging at 100×. **B)** Two-dimensional surface area of myxospore clumps ≥1000 $\mu m^2$. Samples denoted with (i) an asterisk (*****), (ii) star (★), and (iii) diamond (◆) displayed statistically significant ($p \leq 0.05$) differences in distribution relative to (i) WT myxospores without sonication, (ii) WT myxospores with sonication, and (iii) each other, respectively, as determined via two-tailed Mann-Whitney U-tests.

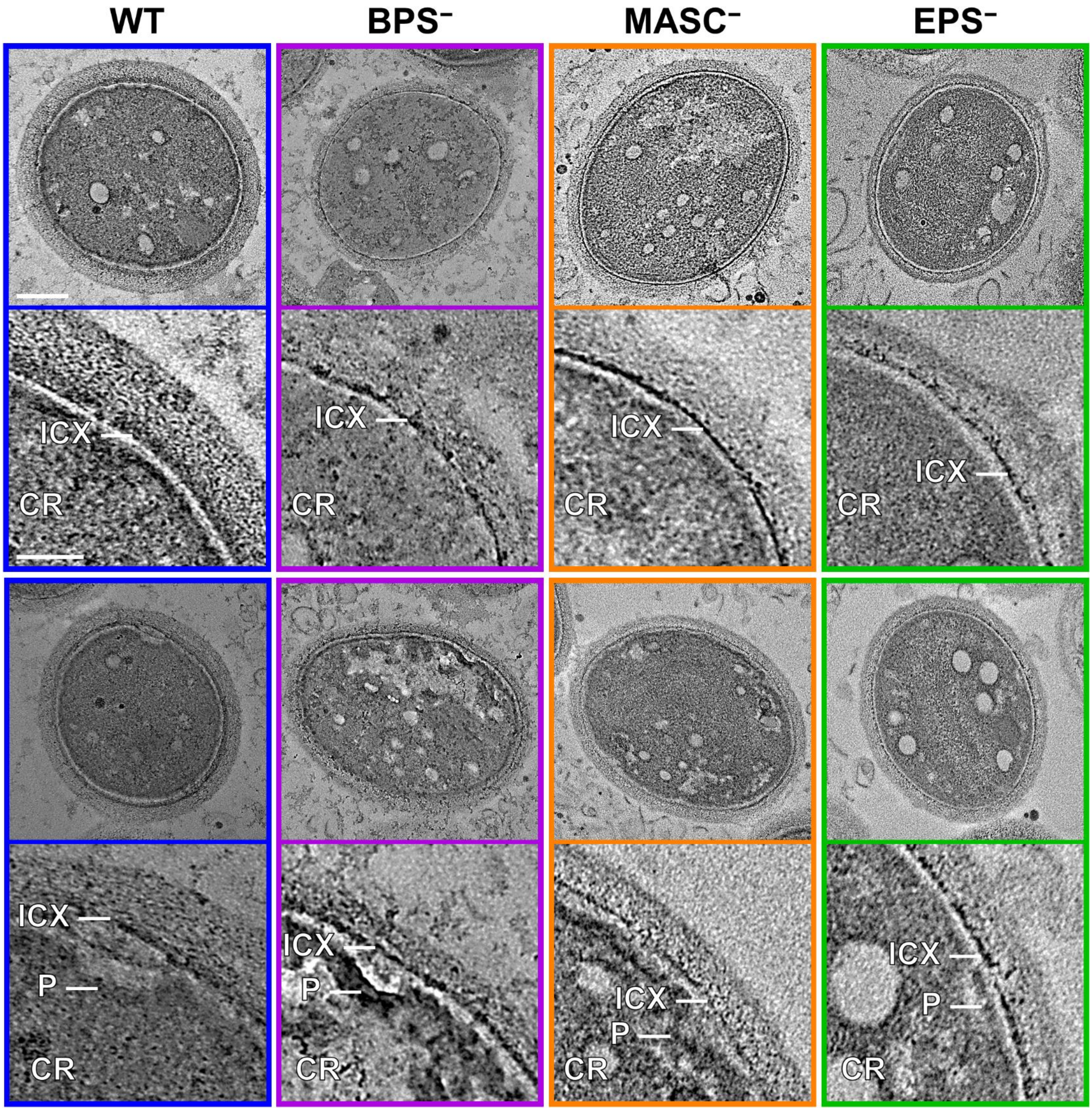

**Supplementary Figure 3. Transmission electron microscopy imaging of natural myxospore envelope architecture in the absence of heavy-metal staining.** Myxospores were obtained by scraping 5-day-old fruiting bodies and immediately immersing them in fixative solution. *Upper panels:* Whole-myxospore view. Scale bar = 300 µm. *Lower panels:* Magnified view of top-right quadrant from upper panels. Scale bar = 100 µm). CR, core; ICX, inner cortex. A redundant set of images has been provided in the lower half of the figure.

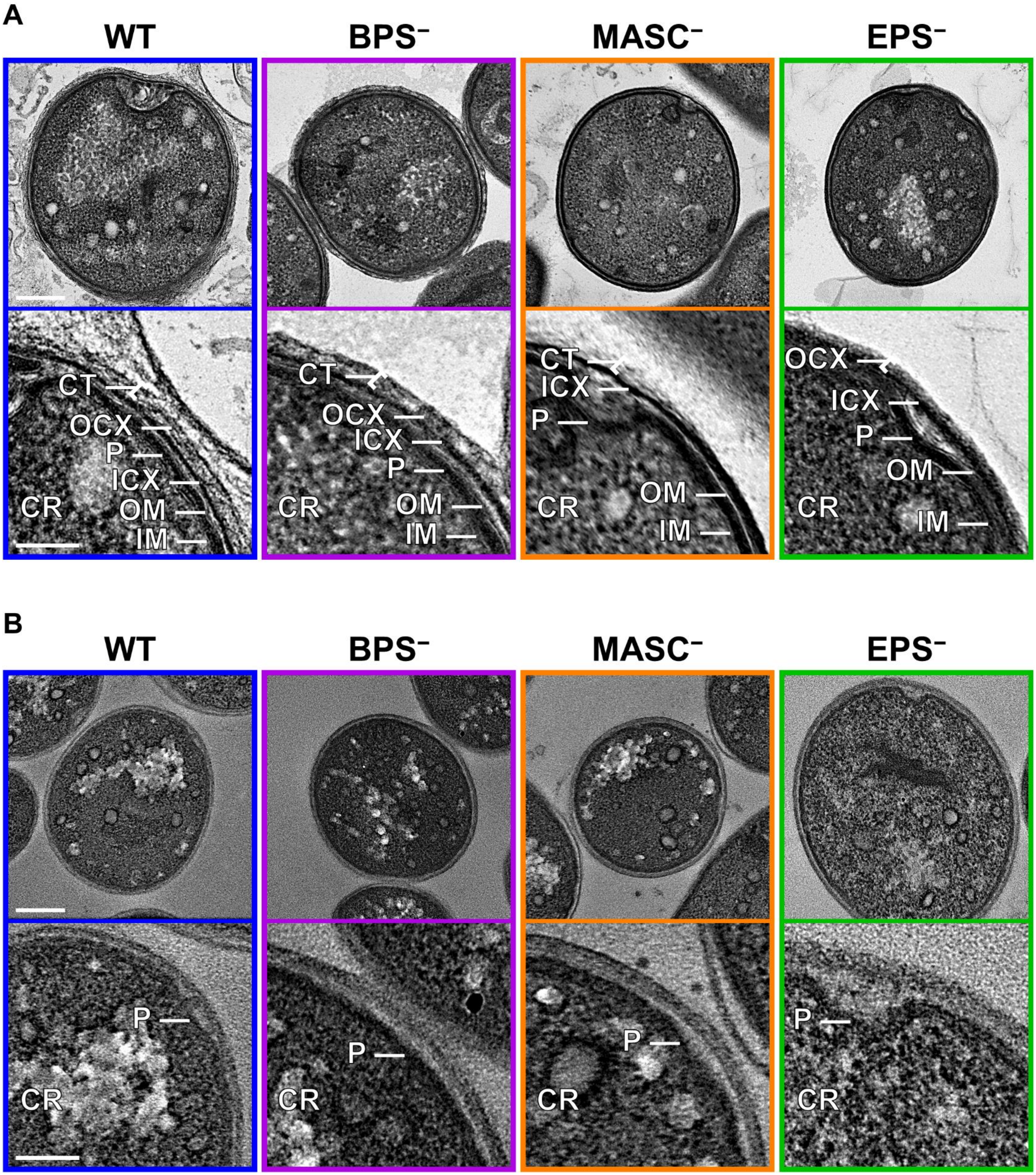
A
WT
BPS⁻
MASC⁻
EPS⁻
CT
OCX
P
ICX
OM
IM
CR
B
P

**Supplementary Figure 4. Transmission electron microscopy imaging of glycerol-induced-myxospore envelope architecture.** Labelling was carried out with $OsO_4$ and tannic acid treatment **(A)** in the presence and **(B)** absence of uranyl acetate and lead citrate staining. *Upper panels:* Whole-myxospore view. Scale bar = 300 μm. *Lower panels:* Magnified view of top-right quadrant from upper panels. Scale bar = 100 μm). CR, core; IM, inner membrane; P, periplasm; OM, outer membrane; ICX, inner cortex; OCX, outer cortex; CT, coat.

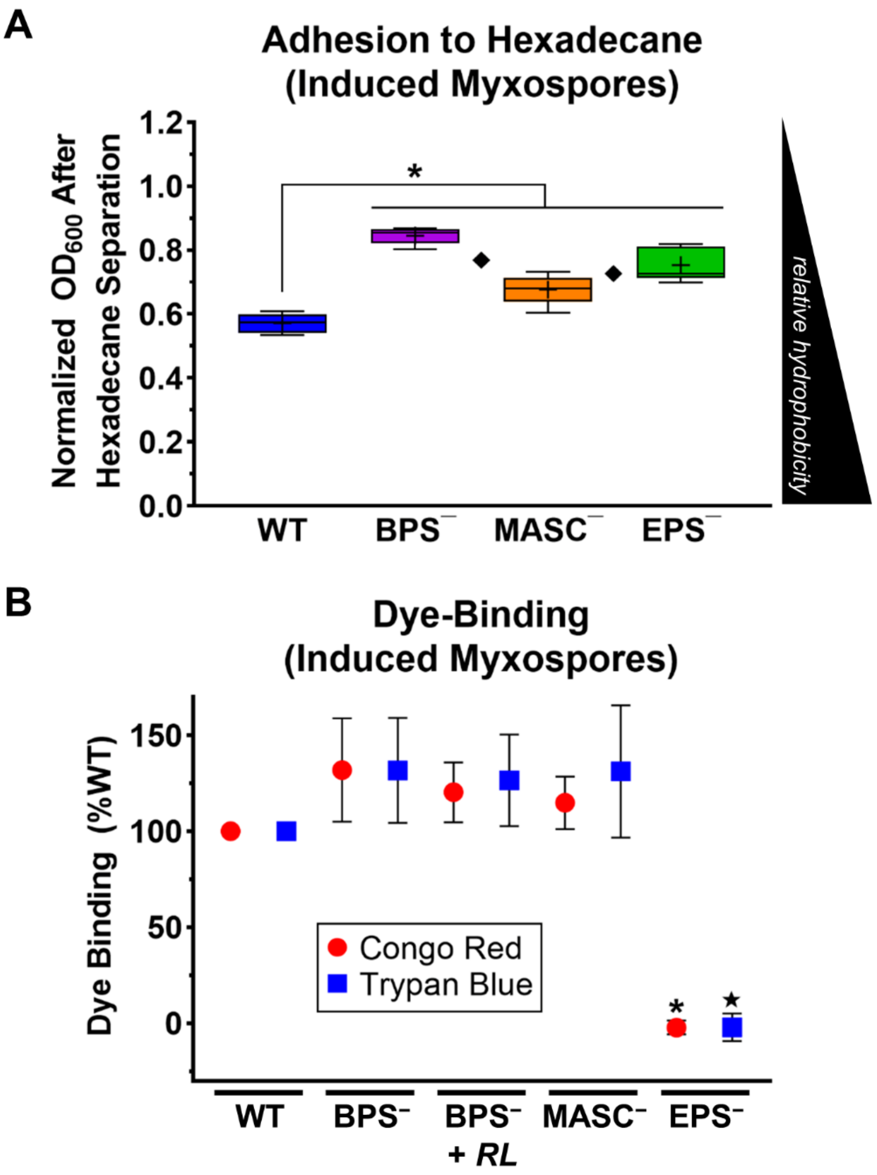

**Supplementary Figure 5. Surface properties of glycerol-induced myxospores. A)** $OD_{600}$ readings of cuvettes containing myxospores in aqueous buffer mixed with hexadecane, after 1 h of emulsion separation. Lower and upper box limits denote $25^{th}$ and $75^{th}$ percentile values, with the lower/upper Tukey fences, median (solid line), and mean (+) values indicated. Mutant myxospore samples denoted with (i) an asterisk (*****) and (ii) a diamond (◆) displayed statistically significant differences in distribution relative to (i) WT myxospores ($p \leq 0.0087$) and (ii) each other ($p \leq 0.0303$), respectively, as determined via two-tailed Mann-Whitney U-tests. For WT, $BPS^{-}$, $MASC^{-}$, and $EPS^{-}$, n = 6, 6, 5, 7, respectively. **B)** Mean values for Congo Red and Trypan Blue dye binding relative to WT myxospores (n = 3), displayed ± standard error of the mean. Mutant myxospore samples denoted with (i) an asterisk (*****) and (ii) star (★) displayed statistically significant ($p \leq 0.0001$) differences in mean values compared to the respective WT reference, as determined via two-tailed Student's t-tests.

**Supplementary Table 1: *Myxococcus xanthus* strains & plasmids used in this study**

| **Strain Code** | **Strain** | **Construction** | **Source** |
|---|---|---|---|
| DZ2 | Wild type | – | [1] |
| TM469 | Δ*wzaX* (i.e. Δ*mxan_7417/epsY*)<br>Note: EPS⁻ strain | – | [2] |
| TM484 | Δ*wzaS* (i.e. Δ*mxan_3225/exoA/fdgA*)<br>Note: MASC⁻ strain | – | [2] |
| TM529 | Δ*wzaB* (i.e. Δ*mxan_1915*)<br>Note: BPS⁻ strain | – | [2] |
| SI130 | Δ*wzaX* IMss-mCherry | TM469 + pSWU19-$P_{pilA}$-IMss-*mCherry* | This study |
| **Plasmid Name** | **Description** | **Construction** | **Source** |
| pSWU19-$P_{pilA}$-IMss-*mCherry* | Red-fluorescent mCherry encoded with an N-terminal signal sequence to target it to the periplasmic leaflet of the inner membrane. Construct is under constitutive expression of the *pilA* promoter | – | [3] |

[1] Campos JM, Zusman DR. Regulation of development in *Myxococcus xanthus*: effect of 3':5'-cyclic AMP, ADP, and nutrition. *Proc. Natl. Acad. Sci. U. S. A.* 1975. 72:518-522.

[2] Ducret A, Valignat M-P, Mouhamar F, Mignot T, Theodoly O. Wet-surface-enhanced ellipsometric contrast microscopy identifies slime as a major adhesion factor during bacterial surface motility. *Proc. Natl. Acad. Sci. U. S. A.* 2012. 109:10036-10041.

[3] Ducret A, Fleuchot B, Bergam P, Mignot T. Direct live imaging of cell–cell protein transfer by transient outer membrane fusion in *Myxococcus xanthus*. *eLife* 2013. 2:e00868.
